# Does Vision Extract Absolute Distance from Vergence?

**DOI:** 10.1101/731109

**Authors:** Paul Linton

## Abstract

Since Kepler (1604) and Descartes (1638), ‘vergence’ (the angular rotation of the eyes) has been thought of as one of our most important absolute distance cues. But vergence has never been tested as an absolute distance cue divorced from obvious confounding cues such as binocular disparity. In this article we control for these confounding cues for the first time by gradually manipulating vergence, and find that observers fail to accurately judge distance from vergence. We consider a number of different interpretations of these results, and argue that the most principled response to these results is to question the general effectiveness of vergence as an absolute distance cue. Given other absolute distance cues (such as motion parallax and vertical disparities) are limited in application, this poses a real challenge to our contemporary understanding of visual scale.

## 1. Introduction

The closer the object is, the more the two eyes have to rotate to fixate upon it. Since Kepler (1604) and Descartes (1637) this mechanism, known as ‘vergence’, has been thought of as one of the visual system’s most important absolute distance cues. This is for eight reasons:

1. Triangulation: Extracting absolute distance from vergence relies on simple principles of geometry. There is no need to infer 3D content from the 2D retinal images. Instead, the visual system is able to triangulate distance from the rotation of the eyes (Parker, Smith, & Krug, 2016; Banks, Hoffman, Kim, & Wetzstein, 2016; Wolfe et al., 2019). Theoretically, this is true of other cues to absolute distance as well (such as accommodation and motion parallax) which is why “conventional wisdom” has traditionally identified “eye vergence, accommodation (focusing the image), binocular disparity, and motion parallax” as the four “primary cues” to depth (Rogers, 2017). See Bishop & Pettigrew (1986) for an optimistic account of 3D vision without the need to infer 3D content from the 2D retinal images, and Clark & Yuille (1990), Ch.1 for a skeptical one.

2. Computer Vision: If you were to reverse engineer distance estimates for a visual system based on two rotating cameras (or eyes), vergence would seems like the natural solution. Indeed, vergence played a central role in the ‘active vision’ revolution in computer vision in the late 1980s and early 1990s (see Krotkov & Kories, 1988; Krotkov, Fuma, & Summers, 1988; Abbott & Ahuja, 1988; Geiger & Yuille, 1989; Krotkov, 1989; Krotkov, Henriksen, & Kories, 1990; Abbott & Ahuja, 1990; Olson & Coombs, 1991; Blake & Yuille, 1992, esp. Ch.8: Brown et al., 1992; Coombs & Brown, 1992; Coombs & Brown, 1993; Krotkov & Bajcsy, 1993; see also Schechner & Kiryati, 2000 for an influential discussion of distance triangulation in computer vision).

3. Effectiveness: Vergence is thought to be particularly effective compared to other absolute distance cues, such as accommodation and motion parallax. In their systematic review of the literature Thompson, Fleming, Creem-Regehr, & Stefanucci (2011) identify just four key absolute distance cues: accommodation, vergence, height in the visual scene, and familiar size. They leave a ‘?’ next to motion parallax. (An evaluation they confirm and further justify in Creem-Regehr, Stefanucci, & Thompson, 2015). Similarly vergence is the principal absolute distance cue discussed by Vishwanath (2014; 2019) and Rogers (2019) in their recent debate on 3D vision, with Rogers (2019) asserting: “No one would deny that binocular disparities and eye vergence are sufficient to ‘specify perceived depth relations’”. Indeed, Rogers (2019) identifies just two cues to absolute distance: vergence and vertical disparities.

Compare the effectiveness of vergence as an absolute distance cue, with the effectiveness of accommodation, motion parallax, vertical disparities, and familiar size:

Accommodation: Mon-Williams & Tresilian (2000) found the variance in pointing responses based on accommodation to be so great that they concluded that “it is clear that accommodation is providing no functionally useful metric distance information for these observers. The responses were unrelated to the actual distance of the target.”

Motion Parallax: Motion parallax has proved a largely ineffective size and distance cue in virtual reality (Beall, Loomis, Philbeck, & Fikes, 1995; Luo, Kenyon, Kamper, Sandin, & DeFanti, 2007; Jones, Swan, Singh, Kolstad, & Ellis, 2008; Jones, Swan, Singh, & Ellis, 2011; Luo, Kenyon, Kamper, Sandin, & DeFanti, 2015), leading Renner, Velichkovsky, & Helmert (2013) to conclude that “there is no empirical evidence that providing motion parallax improves distance perception in virtual environments.” It is for similar reasons that Thompson et al. (2011) and Creem-Regehr, Stefanucci, & Thompson (2015) leave a ‘?’ next to motion parallax as an absolute distance cue. Similarly, Rogers (2019) regards motion parallax as a merely relative depth cue.

Vertical Disparities: To our knowledge there has only been one experiment that indicates that vertical disparities are an absolute distance cue for an object viewed in isolation: Appendix A of Rogers & Bradshaw (1995). This paper lays down very specific criteria for vertical disparity’s effectiveness: the object has to (a) be a fronto-parallel surface, that is (b) covered in a regular texture, and that (c) takes up at least 20° of the visual field. Anything less than 20°, and Rogers & Bradshaw (1995) find that vergence determines absolute distance. Since we rarely encounter objects that take up 20° of the visual field, this cue is very limited in application.

Familiar Size: In a series of papers over 30 years, Walter Gogel (Gogel, 1969; Gogel, 1976; Gogel & Da Silva, 1987; Gogel, 1998) and John Predebon (Predebon, 1979; Predebon, 1987; Predebon, 1990; Predebon, 1992a; Predebon, 1992b; Predebon, 1993; Predebon, 1994; Predebon & Woolley, 1994) questioned whether familiar size really affects our visual perception of scale, and found (in the words of Predebon, 1992b) that “the influence of familiar size on estimates of size mainly reflects the intrusion of nonperceptual processes in spatial responses.” Citing this literature Vishwanath (2014) concludes: “There are no studies that have conclusively demonstrated that familiar size is an independent quantitative perceptual cue to distance. The most recent consensus is that, on its own, familiar size only affects the cognitive inference of distance (Gogel & Da Silva, 1987; Predebon, 1993).”

Vergence: By contrast, Cutting & Vishton (1995) (the standard reference in contemporary textbooks; see Goldstein & Brockmole, 2016; Thompson et al., 2011) suggest that oculomotor cues “could be extremely effective in measuring distance, yielding metric information within near distance”. Empirical evidence for this claim dates back to Meyer (1842), Wheatstone (1852), and Baird (1903). Swenson (1932) found a relationship of *y* = *x* – 0.15 with hidden-hand pointing for distances between 25cm to 40cm, with an average error of 0.17cm, whilst Von Hofsten (1976) found a relationship of *y* = 0.9*x* + 8.5 for distances between 60cm and 118cm, with an average error of 2.2cm. Foley (1980) analysed a series of binocular depth distortions (depth constancy in binocular stereopsis, curvature of the fronto-parallel plane, inability to bisect distances) and argued that they all originated from the same misestimation of distance from vergence. Whilst Foley (1980) helped to cement vergence as an effective absolute distance cue, it also implied that vergence was ‘non-veridical’ (with the visual system’s estimate of the vergence angle being only half its true value), and became the received wisdom for the next two decades.

Foley (1980) set the tone for more recent debates, starting with Mon-Williams & Tresilian (1999), where there is no question vergence is an effective absolute distance cue. The only question is whether vergence is veridical or not? Mon-Williams & Tresilian (1999) is the most influential study on vergence as an absolute distance cue. They found a strong linear relationship between vergence and hidden-hand pointing to the distance of a point of light of *y* = 0.86*x* + 6.5 for distances between 20cm and 60cm (see Fig.1). In this paper, and in Mon-Williams, Tresilian, McIntosh, & Milner (2001), they therefore challenge Foley (1980)’s contention that vergence is a non-veridical distance cue. Instead, they suggest that any compression in their results (a gain of 0.86, rather than a gain of 1) is due to a cognitive strategy that subjects adopt to slightly hedge their bets towards the mean (Poulton, 1980; Poulton, 1988).

**Figure 1.**
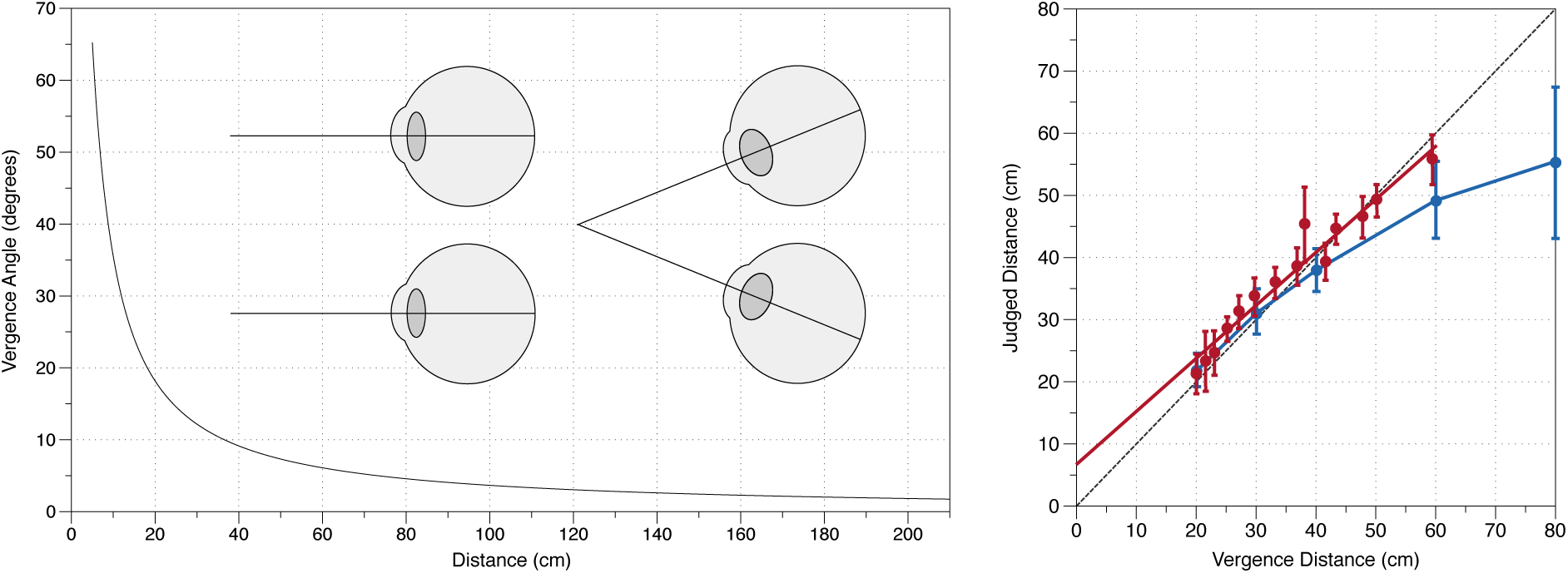
Distance from Vergence. Left panel illustrates how the vergence angle changes with fixation distance. Right panel illustrates the results of absolute distance from vergence in Mon-Williams & Tresilian (1999) (in red), and Viguier, Clément, & Trotter (2001) (in blue), compared to veridical performance (the black dotted line).

Viguier, Clément, & Trotter (2001) is another influential study. They presented subjects with a 0.57° disc at distances between 20cm and 80cm for 5s, and then after 5s in darkness asked subjects to match the distance with a visible reference. They found subjects were close to veridical for 20cm, 30cm, and 40cm, but distances were increasingly underestimated beyond that (60cm was judged to be 50cm, and 80cm was judged to be 56cm; see Fig.1). They conclude that:

> “…in agreement with previous studies (Von Hofsten 1976; Foley 1980; Brenner & van Damme 1998; Mon-Williams & Tresilian 1999; Tresilian et al. 1999) the results of our experiment indicate that vergence can be used to reliably evaluate target distance. This is particularly effective in the near visual space corresponding to arm’s length.”

Whilst Viguier et al. (2001) confirms the effectiveness of vergence as an absolute distance cue, the underestimation of distances beyond 40cm appears to challenge the suggestion that vergence is veridical. In response, Scarfe & Hibbard (2017) ask whether even this underestimation can “in some senses be considered optimal?” Mon-Williams & Tresilian (1999) and Tresilian et al. (1999) observe that symmetric angular noise in the vergence signal will skew the range of probable distances asymmetrically towards further distances. Scarfe & Hibbard (2017) show that although the range of probable distances is skewed towards further distances, the most likely distance actually reduces, explaining the underestimation of distance we observe in Viguier et al. (2001).

However, there is a much simpler explanation. Viguier et al. (2001) use an extended stimulus with a constant angular size, which acts as a counter-cue to vergence. Since a drop-off in performance is not observed after 40cm in Mon-Williams & Tresilian (1999) when a dot (with no discriminable angular size) is used, this appears to be the most likely explanation. So arguably Mon-Williams & Tresilian (1999)’s suggestion that vergence is a veridical cue still stands.

4. Peripersonal vs Extrapersonal Space: Because vergence falls off with the tangent of the distance (Fig.1), there is little change in the vergence angle beyond 2m, and it is commonly suggested that vergence’s effective range doesn’t extend much beyond this (Collewijn & Erkelens, 1990; Cutting & Vishton, 1995; Mon-Williams & Tresilian, 1999; Howard, 2012). One exception is Rogers (2019) who suggests: “The vergence signal indicates viewing at a large distance … signaling that the objects in the scene are (and are seen to be at) a large distance away” (see also Brenner & van Damme, 1998’s suggestion that vergence can scale the distance of a bird in the sky). Two other notable vision scientists have also suggested to me that vergence may be effective for far distances. One reason for believing this is that disparity scaling is effective beyond 2m, so the distance information required for disparity scaling (vergence) must be effective beyond 2m.

But the importance of vergence as an absolute distance cue is especially apparent if we believe that there is an important distinction between near (peripersonal) space and far (extrapersonal) space. This segmentation of visual space was a defining feature of Cutting & Vishton (1995)’s influential review of absolute distance cues, and continues to influence the debate with the suggestion that “vergence of the eyes may provide a key signal for encoding near space” (Culham, Gallivan, Cavina-Pratesi, & Quinlan, 2008), and Creem-Regehr et al. (2015):

> “Binocular stereo provides accurate absolute distance information only in personal space, where it functions to support reaching. Eye-height-scaled perspective is ineffective in both personal space and vista space, but can support accurately scaled egocentric distance judgments in action space, where it helps to control locomotion.”

5. Reaching & Grasping: Vergence is regarded as the preeminent absolute distance cue for reaching and grasping. Bradshaw et al. (2004) find that “vergence information dominates the control of the transport [reaching] component with minimal contribution from pictorial cues”, and suggest that their results confirm Mon-Williams & Dijkerman (1999). Mon-Williams & Dijkerman (1999), Mon-Williams et al. (2001), and Melmoth, Storoni, Todd, Finlay, & Grant (2007), used prisms to manipulate vergence and demonstrate its effect on reaching. Mon-Williams et al. (2001) find that patient DF’s (visual form agnosia) pointing responses almost perfectly mapped the vergence manipulation (*y* = 1.00*x* + 2.8 for base-in prism, *y* = 0.99*x* + 0.22 for no prism, and *y* = 0.96*x* + 0.6 for base-out prism). Culham et al. (2008) also cite earlier behavioural studies that “suggest that eye position and vergence play an important role in the accuracy of reaching movements (Bock, 1986; Henriques & Crawford, 2000; Henriques, Klier, Smith, Lowy, & Crawford, 1998; Henriques, Medendorp, Gielen, & Crawford, 2003; Neggers & Bekkering, 1999; van Donkelaar & Staub, 2000).” Other recent studies that either explore or assume the pre-eminence of vergence as an absolute distance cue for reaching and grasping include: Naceri, Chellali, & Hoinville (2011) (who found results similar to Viguier et al., 2001 for reaching and grasping a fixed-angular sized object in virtual reality); Naceri, Moscatelli, & Chellali (2015); Klinghammer, Schütz, Blohm, & Fiehler (2016); Campagnoli, Croom, & Domini (2017); Grant & Conway (2019); Campagnoli & Domini (2019).

6. Brain Imaging: Brain imaging studies also suggest that vergence acts as the primary absolute distance cue for reaching. Quinlan & Culham (2007) found that the dorsal parieto-occipital sulcus (dPOS) demonstrates a near-space preference, with high activation at closer distances. Importantly, they found that this activation arose when oculomotor cues (vergence, accommodation) were the only cues to absolute distance, leading Quinlan & Culham (2007) to conclude that “it appears that humans do have a functional area that can reflect object distance based on oculomotor cues alone.” This finding was significant for another reason, namely that the same region, the superior parieto-occipital cortex (sPOC) is “primarily – if not, exclusively – concerned with the automatic encoding of target information needed for planning the reach (Pisella et al., 2000; Gallivan et al., 2009; Lindner et al., 2010; Vesia et al., 2010; Glover et al., 2012)” (Grant & Conway, 2019; see Culham et al., 2008 for earlier literature), leading Quinlan & Culham (2007) to conclude:

> “To summarize, in the context of earlier literature, our findings suggest that near vergence is coded in dPOS, a region within the dorsal pathway that plays a critical role in reaching, particularly when the target is off-fixation. Eye position signals related to the current degree of vergence in dPOS likely supply the dorsal stream with critically important information about object distance with respect to current gaze.”

This work complements single-cell recordings that identified vergence coding in LGN (Richards, 1968), the visual cortex (Trotter, Celebrini, Stricanne, Thorpe, & Imbert, 1992; Trotter, Stricanne, Celebrini, Thorpe, & Imbert, 1993; Trotter, Celebrini, Stricanne, Thorpe, & Imbert, 1996; Masson, Busettini, & Miles, 1997; Dobbins, Jeo, Fiser, & Allman, 1998; Trotter & Celebrini, 1999), and the parietal cortex (Gnadt & Mays, 1995). These studies were inspired by the fact that the “psychophysical data suggest an important role for vergence” (Trotter et al., 1992); something that Trotter himself confirmed in Viguier et al. (2001), illustrating the important interplay between the neural and psychophysical data on this topic (see also Lehky, Pouget, & Sejnowski, 1990 for an early neural network model of vergence scaling that Trotter et al. 1992 complements).

7. Size Constancy: We cannot divorce the importance of vergence as an absolute distance cue from the central role vergence is supposed to play in scaling the size (size constancy) and 3D shape (depth constancy) of objects. On size constancy, Combe & Wexler (2010) refer to “the common notion that size constancy emerges as a result of retinal and vergence processing alone” (although they suggest that motion parallax can also have a role to play). The role of vergence in size constancy is particularly acute in the Taylor illusion (scaling an after-image of the subject’s hand as the hand is moved forward and backwards in complete darkness), with Taylor (1941) and Mon-Williams, Tresilian, Plooy, Wann, & Broerse (1997) arguing that vergence is solely responsible for the illusion, and Ramsay, Carey, & Jackson (2007) and Sperandio, Kaderali, Chouinard, Frey, & Goodale (2013) only weakly qualifying that conclusion, with Ramsay et al. (2007) observing: “Of course, vergence provides an extremely powerful distance cue”, whilst Sperandio et al. (2013) find that “perceived size changes mainly as a function of the vergence angle of the eyes, underscoring its importance in size-distance scaling.”

8. Depth Constancy: Vergence is also thought to play a central role in the scaling of 3D shape. As Thompson et al. (2016) note, “disparity is usually considered a relative depth cue, distinct from vergence.” As we have already discussed in the context of Rogers & Bradshaw (1995), when the size of the object is less than 20° the scaling of binocular disparity is dominated by vergence rather than by vertical disparities.

Summarising the contemporary literature, then, there seems to be little question that vergence is one of our most important absolute distance cues for near distances. The consensus seems to be that “as targets get nearer, vergence information plays an increasingly important role in distance perception”, with vergence providing “critically important information” in reaching and grasping (Quinlan & Culham, 2007). The only remaining question is whether vergence provides us with ‘veridical’ (or in some sense ‘optimal’) absolute distance information within reaching space? (Mon-Williams & Tresilian, 1999; Mon-Williams et al., 2001; Scarfe & Hibbard, 2017).

However, one startling fact is that to the best of our knowledge vergence has never been tested as an absolute distance cue divorced from obvious confounding cues such as binocular disparity. We can therefore have little confidence that vergence is determining the absolute distance in these experiments rather than these confounding cues.

We identify three confounding cues, all of which are introduced by the stimulus presentation in Mon-Williams & Tresilian (1999) and Viguier et al. (2001). Subjects are sat in complete darkness, with their vergence in a resting state, and then the stimulus is suddenly presented, often as close as 20cm. This introduces three confounding cues:

1. Double Vision (Retinal Disparity) Before Vergence: If the observer’s vergence is in a resting state, and a stimulus is presented as close as 20cm, then it is going to be seen as double before the observer makes their vergence eye movement. But we know from Morrison & Whiteside (1984) that diplopia (double vision) can be an effective absolute distance cue. Morrison & Whiteside (1984) found that 90% of performance in estimating the distance of a point of light between 0.5m-9.2m could be attributed to diplopia, since performance was only degraded by 10% when the stimuli were shown for a brief period (0.1-0.2s; too quick for a vergence response). Although characterised as ‘coarse stereopsis’ by Ogle (1953), recent literature has emphasised how diplopia provides a direct perception of depth, rather than merely being a cognitive cue (Ziegler & Hess, 1997; Lugtigheid, Wilcox, Allison, & Howard, 2014).

2. Changing Retinal Image (Motion on the Retina) During Vergence: The second confounding cue is the motion of the target on the retina (as it moves from the retinal periphery to the fovea) during vergence. When the stimulus is an isolated target viewed in darkness (as it was in Mon-Williams & Tresilian, 1999 and Viguier, Clément, & Trotter, 2001), subjects will literally watch the targets in each eye streak towards each other across the visual field. Given plausible assumptions about our vergence resting state (that our vergence is beyond arms reach when our eyes are relaxed), the motion of the target across the visual field could be used to inform subjects about the absolute distance of the target.

3. Conscious Awareness of Eye Movements During Vergence: If subjects have to make a sudden vergence eye movement in a response to a near target, they will be consciously aware of their own eye movements because they will literally feel their eyes rotating. If subjects have little or no other absolute distance information, they are going to attend to these consciously felt muscular sensations and attach a lot of weight to them. But this is not how we judge distances in everyday viewing (cf. Berkeley, 1709 who argued that it is). Instead, the suggestion in the literature is that the visual system unconsciously processes muscle movements that we don’t notice (sub-threshold extraocular muscle proprioception) or eye movement plans we don’t know about (efference copy). Consequently, it is important to focus on sub-threshold vergence eye movements (eye movements that subjects don’t notice) if we are to get a better understanding of how vergence actually contributes to distance perception everyday viewing.

To summarise, the extensive literature on vergence as an absolute distance cue tests vergence in the presence of an obvious confounding cue (binocular disparity), and in a way that is divorced from everyday viewing (conscious awareness of eye movements).

The concern that binocular disparity might actually explain absolute distance from vergence is not a new one. It formed the basis of Hillebrand (1894)’s critique of Wundt (1862). So it is worth pausing to ask why, a century on, this concern has never been addressed. The answer is that we seem to be faced with an intractable dilemma. Vergence eye movements are driven by diplopia. So, in order to drive a change in vergence, we have to introduce the very confounding cue that we ought to be controlling for. The solution we adopt in this article is to introduce sub-threshold changes in disparity in order to drive vergence (disparity visible to the observer’s visual system), whilst keeping diplopia invisible to the observer (disparity subjectively invisible). In order to achieve this solution we have to manipulate the observer’s vergence gradually, leaving us open to the objection that we are varying vergence too gradually. We address this concern in the discussion. But we highlight from the outset that this is a necessary trade-off that we have intentionally made. To test vergence as an absolute distance cue in the presence of above-threshold disparities is not an option, and we can have no confidence in experimental results that test vergence as an absolute distance cue in this way.

### Experiment 1

#### Methods

The purpose of Experiment 1 was to replicate Mon-Williams & Tresilian (1999) by having subjects point to the distance of a dot between 20cm and 50cm, but unlike Mon-Williams & Tresilian (1999) we control for retinal disparity and the conscious awareness of eye movements by gradually changing vergence between trials whilst subjects observed a fixation target. The apparatus consisted of a viewing box similar to Mon-Williams & Tresilian (1999)’s (45cm high, 28cm wide, and 90cm long) (Fig.2). The inside of the box was painted with blackout paint mixed with sand (a standard technique for optical equipment: see Gerd Neumann Jr in bibliography).

**Figure 2.**
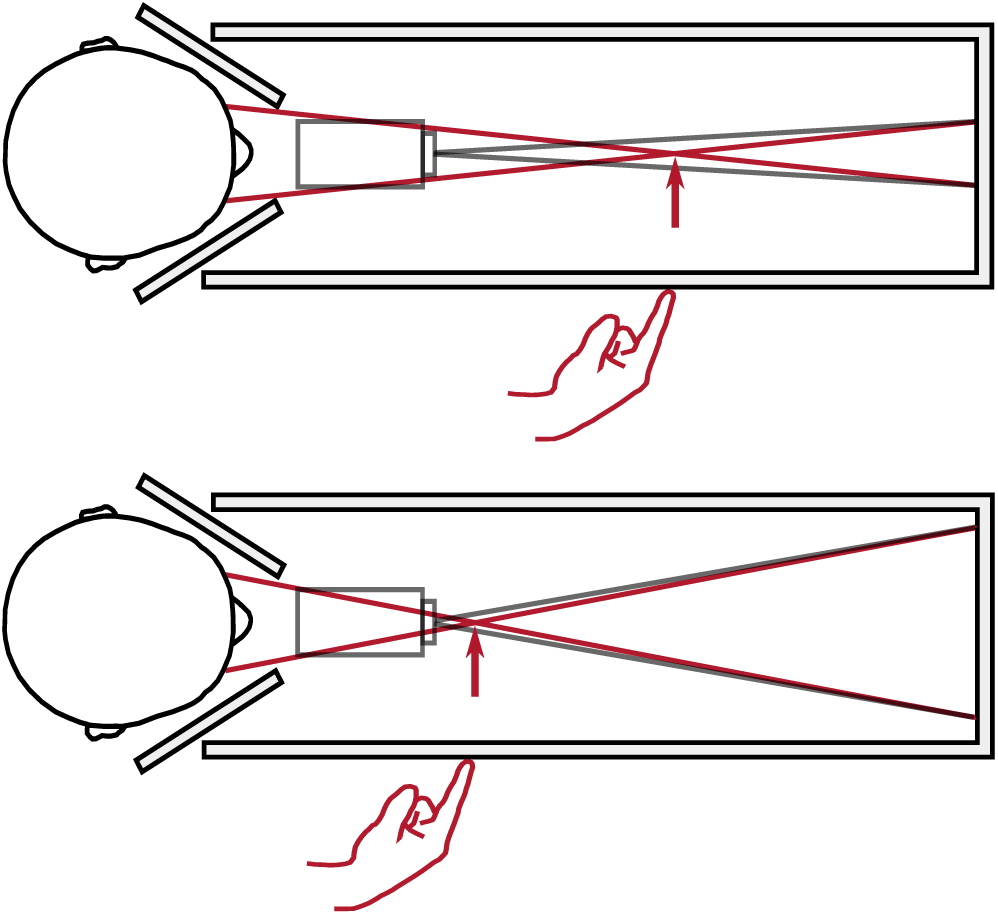
Apparatus for Experiment. Laser projector (in grey) projects two stimuli onto a black metal plate at the end of the apparatus (grey lines). Occluders either side of the head ensure left eye only sees right stimulus, and right eye sees left stimulus (red lines). Vergence specified distance (indicated by red arrow) is manipulated by increasing / decreasing the distance between the two stimuli (compare the upper and lower panels).

Mon-Williams & Tresilian (1999)’s experimental set-up is altered in two fundamental ways:

1. Stimulus alignment: Mon-Williams & Tresilian (1999) align their stimuli with the subject’s right eye, and only vary the vergence demand of the left eye. This approach has two shortcomings: First, it leads to an asymmetric vergence demand. For a 20cm target aligned with the right eye, the vergence demand is 17.75° for the left eye and 0° for the right eye, rather than the symmetric 8.81° for each eye. In normal viewing conditions such extreme asymmetries are eradicated by head rotation. Second, the stimulus is liable to be perceived as drifting rightwards as it gets closer: at 50cm a stimulus aligned with the right eye is offset from the subject’s midline by 3.5°, whilst at 20cm it is offset by 8.8°.

2. Stimulus presentation: We use a laser projector (Sony MP-CL1A), fixed 25cm in front and 5cm below the line of sight, to project the stimuli onto the back wall of the apparatus. In piloting we found that this was the only cost-effective way to ensure that the stimulus was viewed in perfect darkness (lasers emit no light for black pixels, eradicating the residual luminance of CRT, LCD, and LED displays). The stimuli were presented at eye level using PsychoPy (Peirce, 2007; 2009; Peirce et al., 2019). They comprised of (a) a fixation target (300 green dots located within a 1.83° circle, with the dots randomly relocating every 50ms within the circle, creating a shimmering appearance) (Fig.3), and (b) a single green dot that subjects had to point to. The fixation target changed in size sinusoidally between 1.83° and 0.91° at 1Hz. The shimmering appearance and constantly changing size of the fixation target ensured that as the vergence demand was varied, any residual motion-in-depth from retinal slip would be hard to detect.

**Figure 3.**
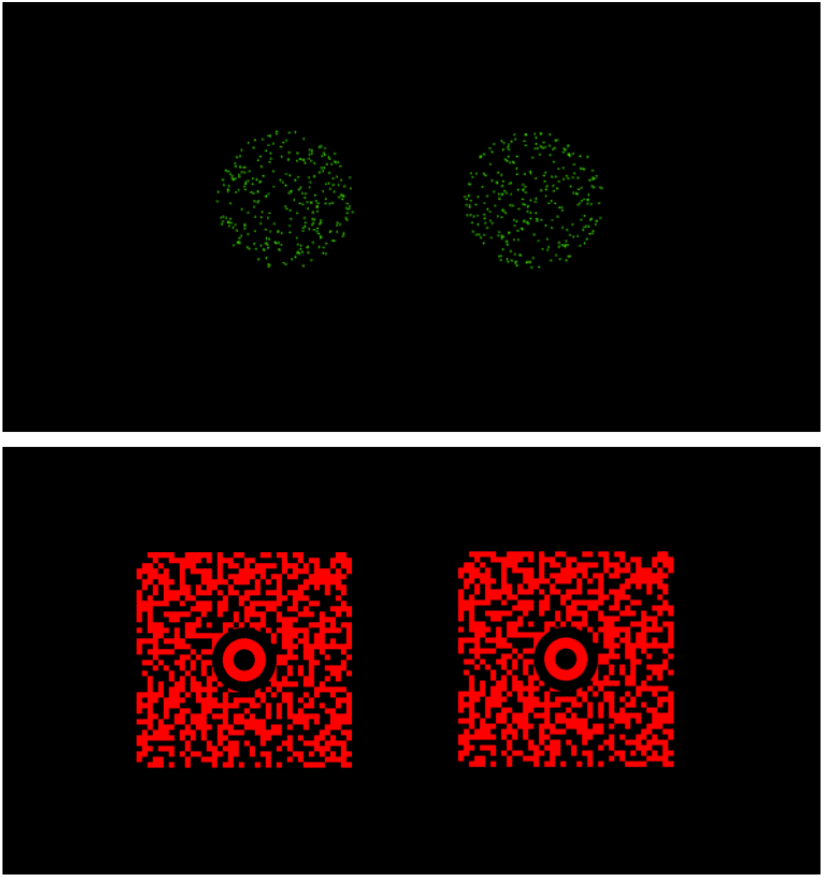
Fixation Targets for Experiment 1 (top) and Experiment 2 (bottom). The fixation target is seen between trials (whilst vergence is gradually changing). Note that the actual stimulus that subjects point to during each trial (once vergence is stationary) is a dot.

We set the initial vergence distance when subjects enter the apparatus at 50cm. During the first trial, which was longer than all the subsequent trials, the vergence distance was varied over 32 seconds from 50cm to 29cm (the centre of the range) as the subjects observed the fixation target. After 32 seconds, a dot was presented at 29cm and subjects had to point to its distance on the side of the box. In each subsequent trial the vergence distance was stepped up or stepped down over 15 seconds (as subjects observed the fixation target) by one step in a pseudo-random walk that covered 7 vergence-specified distances (20cm, 22cm, 25cm, 29cm, 34cm, 40cm, and 50cm), after which a dot was presented, and subjects had to point to the distance of the dot. Each subject completed 96 trials (4 sets of 24 trials), with the pseudo-random walk ensuring that each of the 7 distances was tested at least 10 times.

All subjects were naïve as to the purpose of the experiment and the experimental set-up. They did not see the room or apparatus beforehand, which was sealed off by a curtain, and they were wheeled into the experimental room wearing a blindfold. Their hand was guided to a head and chin rest, and they had to ensure their head was in place, with a further hood of blackout fabric pulled over their head, before they could take the blindfold off. This procedure, coupled with the fact that the box was sealed, and the illumination in the room outside the box was reduced to a low-powered LED, ensured that the stimuli were viewed in perfect darkness, and the subjects had as few prior assumptions about the distances being tested as possible.

To ensure binocular fusion, before each of the 4 sets of trials we had subjects confirm they could see the target monocularly in each eye before opening both eyes. If fusion failed, subjects would experience diplopia. We asked them to inform us immediately if the target was seen as double. That set of trials was immediately paused, and restarted after a break. We also checked with each subject during the break between trials that the stimulus wasn’t seen as double.

After the main experiment was completed, a control study was run in full-cue conditions to confirm Mon-Williams & Tresilian (1999)’s and Swan et al. (2015)’s finding that hidden hand pointing is a good reporting mechanism for perceived distance. The control replicated the head rest, chin rest, and the right-hand wall, of the original apparatus, but removed the top, back, and left-hand wall, enabling a familiar object (a 510g Kellogg’s Rice Krispies box) to be seen in full-cue conditions. Subjects pointed to the front of the cereal box with a hidden hand in 3 sets of trials that ensured 10 trials in total for each of the 7 distances (20cm, 22cm, 25cm, 29cm, 34cm, 40cm, and 50cm). One subject (SM) was unable to return to complete the control.

The observers were 12 acquaintances of the author (9 male, 3 female; ages 28-36, average age 31.2) who indicated their interest to volunteer for the study in response to a Facebook post. Observers either did not need visual correction or wore contact lenses (no glasses). All observers gave their written consent, and the study was approved by the School of Health Sciences Research Ethics Committee, City, University of London in accordance with the Declaration of Helsinki. Three preliminary tests were performed to ensure that (a) the subjects’ arm reach was at least 60cm, (b) their convergence response was within normal bounds (18D or above on a Clement Clarke Intl. horizontal prism bar test), and (c) that their stereoacuity was within normal bounds (60 sec of arc or less on a TNO stereo test).

#### Results

The results for the 12 participants in Experiment 1 are reported in Figure 4.

**Figure 4:**
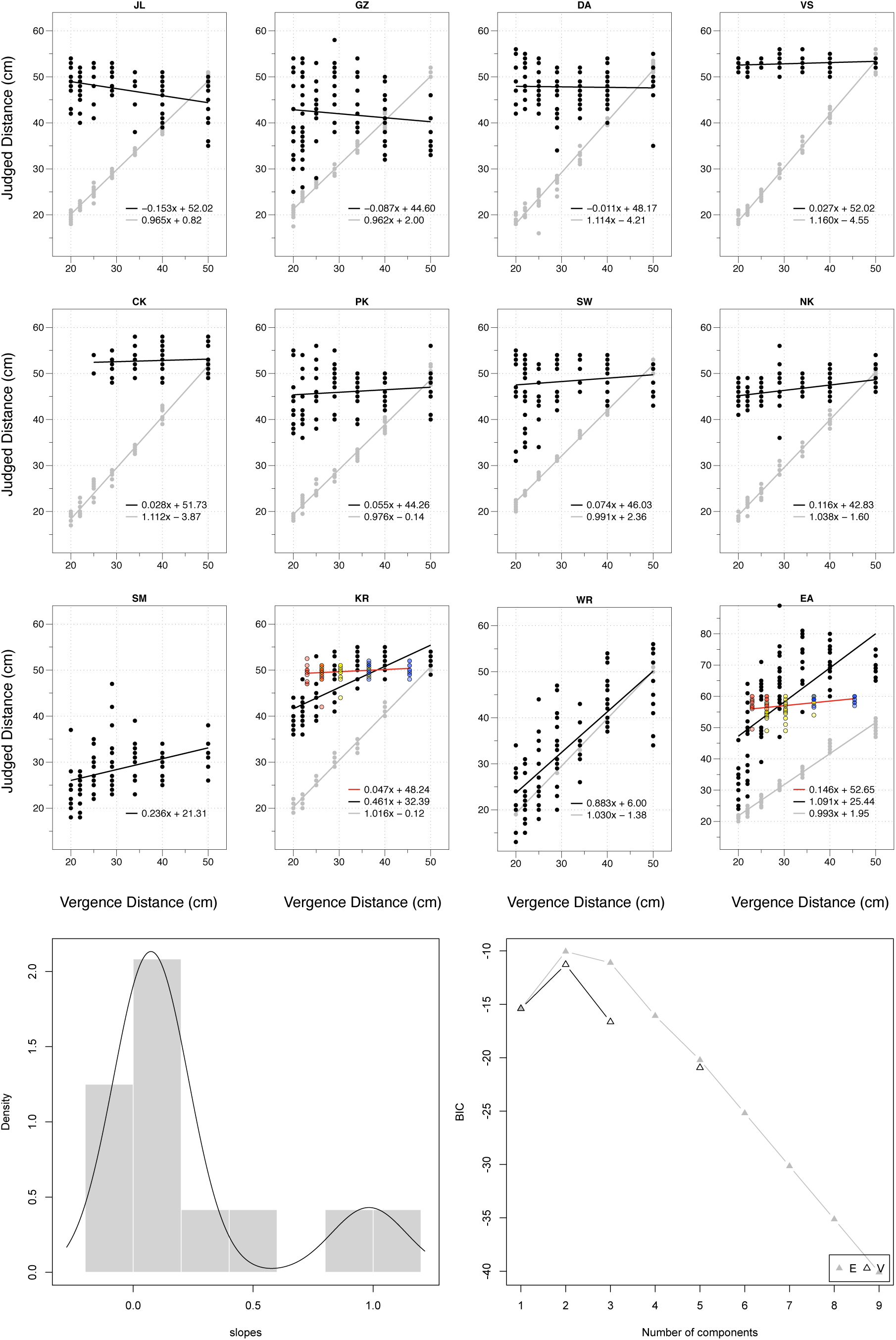
Individual Results for 12 Subjects in Experiment 1. Grey dots and grey line indicate performance in full-cue condition. Black dots and black line indicate performance in vergence-only condition. Coloured dots and red line for subjects KR and EA indicate revised performance in Experiment 2 (the colour of the dots indicates accommodative demand: • = –4.15D, • = – 3.15D, and • = –2.15D). Bottom panels: Results best fit with two populations with equal variance (ΔE), with Gaussian mixture model plotted on histogram of the slopes.

The data were analysed using a linear mixed-effects model (Pinheiro & Bates, 2000) in R (R Core Team, 2012) using the lme4 package (Bates, Maechler, & Bolker, 2012), with pointed distance (*y*) a combination of a fixed-effect of vergence distance (*x*), and a random-effect of each participant (*x* | Subject), so that: *y* ∼ *x* + (*x* | Subject). All data and analysis scripts are accessible in an open access repository: https://osf.io/2xuwn/

As expected, hidden-hand pointing in full-cue condition was close to veridical: *y* = 1.032*x* – 0.76 (with 95% confidence intervals of 0.992 to 1.071 for the slope, and –2.36 to 0.73cm for the intercept). However, it is a different story when vergence is the only cue. We clustered a histogram of the slopes in Fig.4 using the mclust5 package (Scrucca, Fop, Murphy, & Raftery, 2017), and found that the results were best explained by two populations of equal variance:

1. 10 out of the 12 subjects had a slope indistinguishable from 0 (*y* = 0.074*x* + c). Using a linear mixed-effects model we estimated the slope and intercept for these 12 observers to be *y* = 0.075*x* + 43.52 (with 95% confidence intervals of –0.035 to 0.183 for the slope, and 37.12 to 49.70 for the intercept).
2. 2 out of the 12 subjects (EA and WR) had a slope indistinguishable from 1 (*y* = 0.983*x* + c). Using a linear mixed-effects model we estimated the slope and intercept for these 2 observers to be *y* = 0.987*x* + 15.72 (with 95% confidence intervals of 0.747 to 1.219 for the slope, and –3.94 to 36.13 for the intercept).

These results illustrate that vergence was an ineffective absolute distance cue for the vast majority (10 out of 12) of our subjects. Does this, however, mean that vergence was an effective absolute distance cue for our other two subjects, WR and EA? We suggest not. Both WR and EA, along with KR (the subject with the 3rd highest slope), reported symptoms consistent with vergence-accommodation conflict in their debrief. EA and KR reported using the size change of the dot with defocus as a cue to distance, whilst WR complained of significant eye strain, describing the experiment as “exhausting” for his eyes (and abandoned a revised version of the experiment as being “painful” and “quite exhausting for his eyes” after just six trials).

We hypothesised that the performance of these three subjects relied on vergence-accommodation conflict, rather than vergence being an effective absolute distance cue. The purpose of Experiment 2 was to test whether their performance would disappear once vergence-accommodation conflict had been controlled for, as well as testing 12 new participants using our revised experimental paradigm that controlled for vergence-accommodation conflict.

### Experiment 2

#### Methods

In Experiment 2, vergence-accommodation conflict was kept within reasonable bounds (within the ‘zone of clear single binocular vision’: Hoffman, Girshick, Akeley, & Banks, 2008) by dividing the trials into Near (23cm to 30cm), Middle (23cm to 45.5cm), and Far (30cm to 45.5cm) trials, and testing each of these vergence distances with different lenses to ensure that:

- Near: Accommodation set at 24cm (–4.15D) to test vergence at 23cm, 26cm, and 30cm.
- Middle: Accommodation set at 32cm (–3.15D) to test vergence at 23cm, 26cm, 30cm, 36.5cm, and 45.5cm.
- Far: Accommodation set at 47cm (–2.15D) to test vergence at 30cm, 36.5cm, and 45.5cm.

The fixation target was also made larger (2.4° x 2.4°) and higher contrast to aid accommodation (Fig.3). Rather than changing in angular size, the fixation target now varied in luminance (between 100% and 50% of its initial luminance, at 2Hz). To increase contrast, stimuli were projected onto a white screen 156cm away from the observer, rather than a black metal plate 90cm away. To ensure this increase in illumination did not also illuminate the apparatus, black fabric was added to ensure a narrow viewing window, and red filters from red-cyan stereo-glasses (blocking ≈100% green light, ≈90% blue light) were added in front of each eye. In a separate preliminary experiment using an autorefractor these filters were found to have no impact on accommodation.

We set the initial vergence distance to 50cm. During the first trial, which was longer than all the subsequent trials, the vergence distance was varied over 50 seconds from 50cm to the centre of the range (26cm for Near trials, 30cm for Middle trials, and 36.5cm for Far trials) as subjects observed the fixation target, before a dot was presented and subjects pointed to its distance. In each subsequent trial the vergence distance was stepped up or stepped down over 30 seconds (as subjects observed the fixation target) by one step in a pseudo-random walk that covered the vergence-specified distances, before a dot was presented and subjects pointed to its distance. The 12 new participants completed 7 sets of 20 trials (2 Near, 3 Middle, and 2 Far) that ensured that each combination of accommodation and vergence was tested at least 10 times. For the subjects from Experiment 1 a reduced version of the experiment was constructed: 4 sets of 20 trials (1 Near, 2 Middle, and 1 Far) with 23cm and 26cm tested in Near; 26cm, 30cm, and 36.5cm tested in Middle; and 36.5cm and 45.5cm tested in Far. Subject WR from Experiment 1 was unable to return, but subjects KR and EA returned to complete this experiment.

The 12 new participants were 12 City, University of London undergraduate students (8 female, 4 male; age range 18-27, average age 20.8) recruited through flyers and Facebook posts. All subjects were naïve as to the purpose of the experiment. The same exclusion criteria as Experiment 1 were applied, with the additional requirement that subjects’ accommodative responses (tested with a RAF near-point rule) were within normal bounds. The study was approved by the School of Health Sciences Research Ethics Committee, City, University of London in accordance with the Declaration of Helsinki, and all subjects gave their written consent. The undergraduate students were paid £10/hr + a £20 completion bonus.

#### Results

The results for the two subjects from Experiment 1 (KR and EA) are reported alongside their Experiment 1 performance in Figure 4. We find a dramatic reduction in their performance once vergence-accommodation conflict has been controlled for: EA’s previous performance of *y* = 1.091*x* + 25.44 drops to *y* = 0.146*x* + 52.65, whilst KR’s previous performance of *y* = 0.461*x* + 32.39 drops to *y* = 0.047*x* + 48.24 (both drops in performance are significant to p < 0.001). This confirms the hypothesis that their performance in Experiment 1 was driven by vergence-accommodation conflict, rather than vergence being an effective absolute distance cue.

The results for the 12 new participants are summarised in Figure 5, with individual results reported in Figure 6.

**Figure 5.**
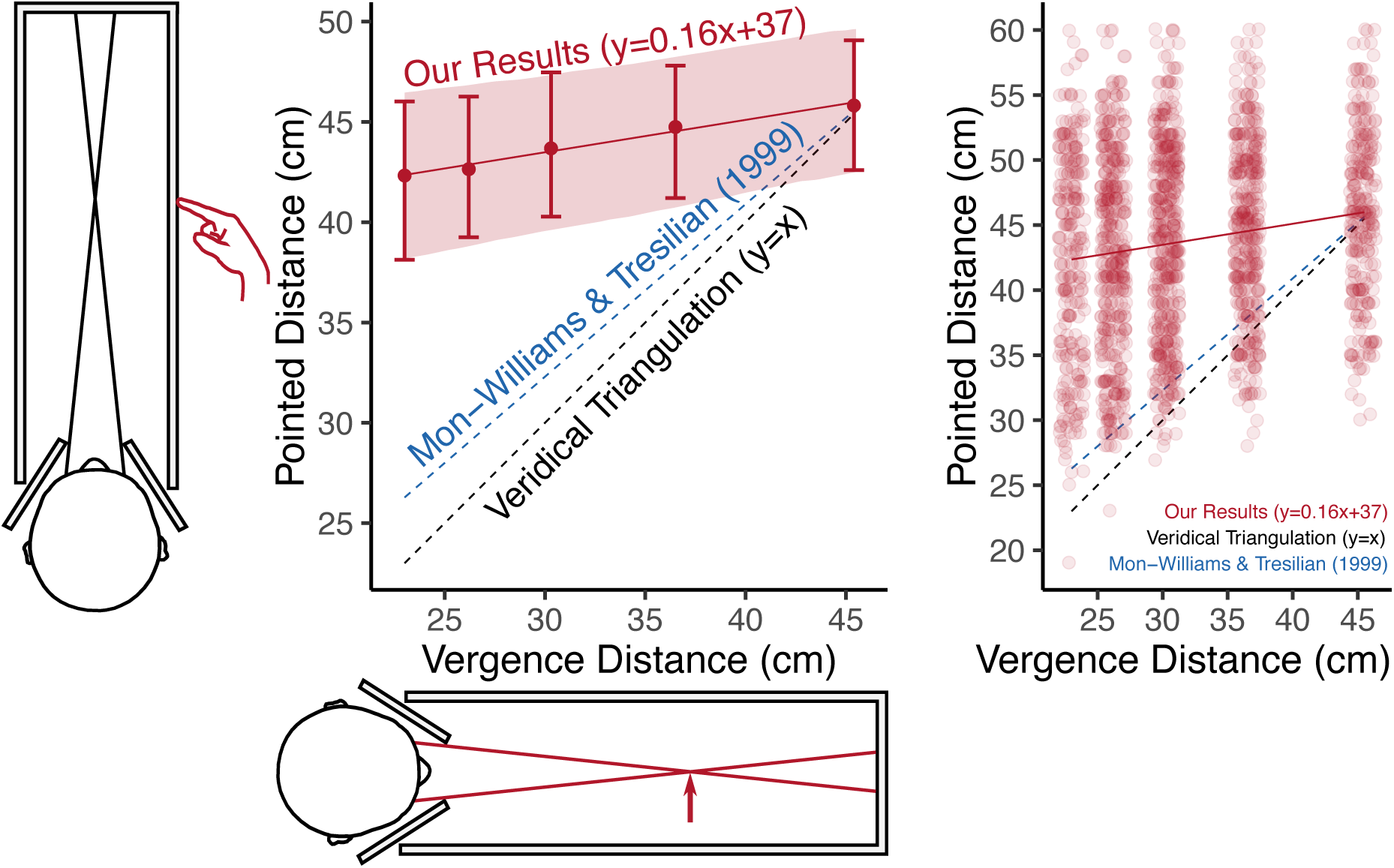
Summary of Results from Experiment 2 when Vergence and Accommodation are the only cues to Distance. On the x-axis is the vergence-specified distance, and on the y-axis the pointed distance. The left panel illustrates averaged results. The error bars represent bootstrapped 95% confidence intervals across observers. The error band represents the bootstrapped 95% confidence interval of the linear mixed-effects model. The right panel plots the raw trial data across observers as a jitter plot.

**Figure 6:**
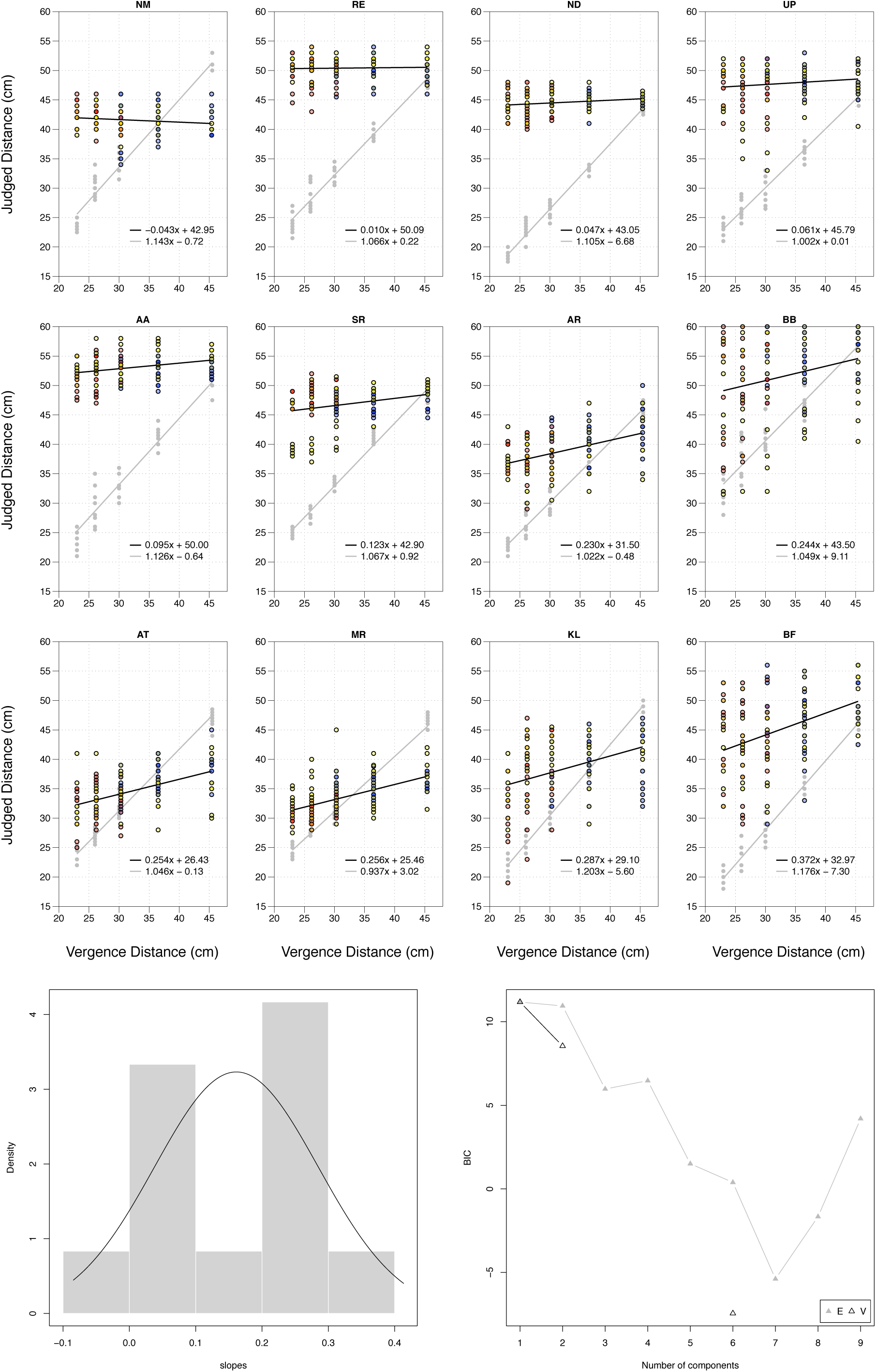
Individual Results for 12 Subjects in Experiment 2. Grey dots and grey line indicate performance in full-cue condition. Coloured dots and black line indicate performance in vergence and accommodation-only condition (colour of the dots indicates accommodative demand: • = – 4.15D, • = –3.15D, and • = –2.15D). Bottom panels: Results best fit with a single population, with Gaussian mixture model plotted on histogram of the slopes.

As expected, the 12 new subjects were close to veridical in full-cue conditions: *y* = 1.078*x* – 0.69 (with 95% confidence intervals of 1.036 to 1.122 for the slope, and –3.19 to 1.81 for the intercept). By contrast, when vergence and accommodation were the only cues to absolute distance, their performance had three defining features:

1. Low Gains: We clustered the individual slopes in Fig.6 using the mclust5 package and found that a single population with an average slope of *y* = 0.161*x* + c best fits the data (although this was only marginally better than two populations with equal variance). Using a linear mixed-effects model we estimated the slope and intercept to be *y* = 0.161*x* + 38.64 (with 95% confidence intervals of 0.090 to 0.239 for the slope, and 33.43 to 43.36cm for the intercept). This confirms similar findings for 11 of the 12 participants in Experiment 1 (when we include KR and EA’s revised results), with the 12^th^ participant unable to return. On the basis of these results we are confident that this is a highly replicable finding.

2. High Variability: The high degree of variance in the results is best illustrated by the raw data of the 12 subjects (Figure 5, right panel). There is little evidence of a ‘specific distance tendency’ (Gogel, 1969). Instead, subjects appear to effectively be guessing.

Indeed, we were struck by the high degree of variance in the results of the 6 subjects with above average slopes. To quantify this variance we estimated the standard deviation of the residual error (i.e. how much each of those 6 subjects departed from their own line of best fit in Fig.6), after correcting for motor error (assuming that perceptual error and motor error are independent). We did this by attributing all of the variance in the full-cue control condition to motor error (an intentional overestimate) in order to produce a conservative estimate of the residual perceptual error using the following formula:

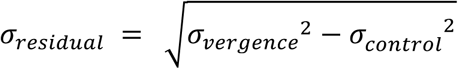

Limiting ourselves to ± 2 standard deviations from the slope of best fit to rule out any outliers, we find an average residual error for those 6 subjects of 17cm. Given the range of the experiment itself was 22.5cm, one is left questioning just how functionally useful an absolute distance cue with this degree of variance could be.

3. No Benefit from Accommodation: It has been shown that accommodation can contribute absolute distance information, although it is subject to a high degree of variance (Fisher & Ciuffreda, 1988; Mon-Williams & Tresilian, 2000). But we found no effect on absolute distance by varying accommodation by two dioptres. We compared (a) the line of best fit when accommodation was varied in line with vergence for 23cm, 30cm, and 45.5cm, to (b) the line of best fit when accommodation was fixed at 30cm for all three distances. We found a slight reduction in performance when accommodation was varied (*y* = 0.147*x* + 38.91) vs. when accommodation was fixed (*y* = 0.176*x* + 38.02), but this effect was not statistically significant. We certainly didn’t find the improvement in performance that one would expect if accommodation were a complementary absolute distance cue to vergence.

## 4. Discussion

These results show that vergence and accommodation were ineffective absolute distance cues for our participants. The distance estimates of some of the participants in Experiment 2 were biased by vergence. But this is not evidence of unconscious processing of the vergence signal. Instead, the subjects with the highest gains reported responding to consciously felt muscular sensations from intense sustained near fixation. For instance BF, the subject with the highest gain, reported that the experiment was “messing up my accommodation”:

> “I could feel my eye are working, my eyes are focusing then relaxing then focusing.”

> “I really had to focus to stop them going two … the target started to separate when I didn’t really focus on it.”

> “I usually get the same sensation when I’m up too late and doing some studies – a slight strain in the eye, it’s not too bad, it’s just that you really have to focus.”

Similarly KL, the subject with the second highest gain, reported that with near targets she felt her “eyes accommodating a lot to get them to work.” So, whilst our experimental paradigm effectively controlled for the conscious muscular sensations that accompany eye movements (kinaesthesia), it failed to control for the conscious muscular sensation of sustained near fixation (proprioception). Such muscular sensations are rarely felt in everyday viewing, and appear to be a shortcoming of manipulating vergence and accommodation as pure optical reflexes (see Charman & Heron, 2015 for similar concerns about Badal systems).

We believe that these results support the conclusion that vergence and accommodation are ineffective absolute distance cues. First, this would be the most parsimonious interpretation of our results. Second, we should be very reluctant to try and construct rationalisations to ‘save the data’ from previous experiments if the data from those experiments were collected in the presence of obvious confounding cues. To repeat, we can have no confidence that those data reflect vergence actually functioning as an absolute distance cue. Third, ‘harking’ (hypothesis after results are known) has little cogent value unless it makes testable predictions (Kerr, 1998). Otherwise, it risks simply becoming a ‘just so’ story to fit the data (Catmur, Press, Cook, Bird, & Heyes, 2014; Heyes, 2019). As Firestone & Scholl (2016) note, albeit in a different context: “We sincerely wish to avoid the specter of vague ‘Australian stepbrothers’ (Bruner & Goodman, 1947…) that merely could explain away these effects, without evidence that they really do.”

However, there are a number of alternative interpretations of these results that we cannot conclusively reject, although we personally find each of them implausible. In the remainder of this Discussion we therefore explore five alternative interpretations of these results that have been put to us. We explain why we do not find these alternative explanations convincing, but we recognise that none of these possibilities can be definitively excluded.

### 1. No Eye-Tracking

We did not use eye-tracking to track the subjects’ vergence, so how do we know that subjects were actually changing their vergence during the experiment?

We did not use eye-tracking for four reasons: First, Hooge, Hessels, & Nyström (2019) found that readily available research eye-trackers “are not accurate enough to be used to determine vergence, distance to the binocular fixation point and fixation disparity”, with errors of up to 2.5°. Second, we share Quinlan & Culham (2007)’s concern that near-infrared light from eye-trackers will stray into the visible spectrum and introduce disparity cues. Third, we were concerned that eye-tracking would be impractical given our use of parallax barriers, making a clear view of both eyes (for the eye-tracker) and the calibration targets (for the observer) impossible. Fourth, we were very careful about the prior knowledge that subjects had about the apparatus, and we feared that calibration would be impossible without compromising this in some way.

Collewijn & Erkelens (1990) are critical of studies that do not provide an objective measure of vergence using eye-tracking. However, we rely on a subjective measure of vergence (diplopia), which we would argue is more reliable than camera-based eye-tracking. Before each set of trials, we asked subjects to confirm they could see the target in each eye monocularly, and then confirm that they could see a single fused target when they opened both eyes. We asked them to report if the target went double at any time during each set of 24 trials (Experiment 1) or 20 trials (Experiment 2). We paused and restarted the experiment after a break if it did. In the break between sets of trials, we also asked the subjects to confirm the target had been fused in the previous set of trials.

Since our target was a single dot, the presence or absence of diplopia provides us with a very effective test of binocular fusion. Schor & Tyler (1981) estimate diplopia thresholds for a fixation dot to be 8 arcmin. Diplopia thresholds for thin vertical bars have been found to be as low as 3 arcmin (Schor & Tyler, 1981) and 5 arcmin (Schor, Wood, & Ogawa, 1984). This helps to explain why nonius lines have traditionally been treated as a gold-standard for vergence “even when”, as Schor, Wood, & Ogawa (1984) note, “small discrepancies between subjective and objective measures of horizontal fixation disparity are taken into account (Kertesz et al., 1983).” Kertesz, Hampton, & Sabrin (1983) found an average diplopia threshold of 6 arcmin for nonius lines, whilst Jaschinski, Bröde, & Griefahn (1999) found diplopia thresholds of 5 arcmin or less when measured binocularly, and 2 arcmin or less when measured with dichoptic nonius lines. In recent work Grove, Finlayson, & Ono (2014) found higher diplopia thresholds (around 13 arcmin for uncrossed disparities and 8 arcmin for crossed disparities), but their vertical bars were 4.4 arcmin wide (vs. 1.5 arcmin dot and 1.5 arcmin lines used by Schor & Tyler, 1981), so their thresholds should arguably be reduced by 3 arcmin to 5-10 arcmin for a dot stimulus.

In conclusion, a best estimate of the accuracy of diplopia thresholds in our experiment should be no more than about 10 arcmin (8 arcmin Schor & Tyler, 1981), and could well be lower if the thin vertical bar / nonius line literature applies. Compare this to objective measures from readily available research eye-trackers, where the 2D gaze literature (Choe, Blake, & Lee, 2016; Drewes, Zhu, Hu, & Hu, 2014; Wildenmann & Schaeffel, 2013; Wyatt, 2010) and the 3D gaze literature (Hooge et al., 2019) report similar errors of magnitude (up to ≈ 2.5°). Since our subjective measures are an order of magnitude (up to 15 times) more accurate than readily available objective measures from eye-tracking (10 arcmin vs 2.5°), we conclude that our subjective test for fusion based on diplopia is to be preferred. We recognise that some authors feel especially strongly that vergence studies should be accompanied by eye-tracking (Collewijn & Erkelens, 1990), but there is reasonable disagreement on this point, and notable studies share our accuracy (Hooge et al., 2019) and logistical (Quinlan & Culham, 2007) concerns.

### 2. Vergence-Accommodation Conflict

Could vergence-accommodation conflict account for our results? We do not believe so for three reasons.

First, as we discussed in the Introduction, there is widespread skepticism that accommodation functions as an effective absolute distance cue. Recall Mon-Williams & Tresilian (2000)’s finding that accommodation provides “no functionally useful metric distance information”.

Second, vergence-accommodation conflict is a facet of most of the studies that demonstrate (close to) veridical absolute distance from vergence. First, any study which varies vergence using prisms, such as Mon-Williams & Tresilian (1999), keeps accommodation fixed. Since Mon-Williams & Tresilian (1999) varied vergence over 3.33D (20-60cm), they induce at least 1.67D of vergence-accommodation conflict, and potentially even more (we were unable to determine the exact figure). Second, any study that relies on a fixed display such as Von Hofsten (1976), is going to induce significant vergence-accommodation conflict. Von Hofsten (1976) found an almost perfect relationship between vergence and perceived distance up to 118cm, at which point there was 1.3D of vergence-accommodation conflict. Third, any study that relies on virtual reality, such as Naceri, Chellali, & Hoinville (2011), is going to induce 4D of vergence-accommodation conflict (their nearest target was 25cm, with accommodation set close to optical infinity), and yet they found results consistent with Viguier et al. (2001). So vergence-accommodation conflict hasn’t previously been an impediment to finding that vergence is an effective absolute distance cue, and the maximum vergence-accommodation conflict within our second experiment (1.17D) is well within the range of these previous experiments.

Third, we explicitly tested the effect of vergence-accommodation conflict in our second experiment by contrasting the results for three vergence distances (23cm, 30cm, and 45.5cm) when (a) there was virtually no vergence-accommodation conflict (accommodation set at: 24cm, 31.5cm, and 46.5cm respectively) vs. (b) when there was up to 1.17D of vergence-accommodation conflict (accommodation set at 31.5cm) and found a non-statistically significant reduction in performance in the no vergence-accommodation conflict condition. Coupled with the fact that subjects KR and EA reduced in performance between Experiment 1 and Experiment 2 (when accommodation cues were provided), and we can conclude that vergence-accommodation conflict is not the explanation.

### 3. Conscious Awareness of Eye Movements

We controlled for conscious awareness of vergence eye movements. One objection is that this is what is meant in the literature by vergence functioning as an effective absolute distance cue. We disagree with this suggestion for five reasons:

First, conscious awareness of our own eye movements is not how we judge distances in everyday viewing. This isn’t what Rogers (2019) means when he suggests: “No one would deny that binocular disparities and eye vergence are sufficient to ‘specify perceived depth relations’”, what Cutting & Vishton (1995) mean when they suggest vergence “could be extremely effective in measuring distance, yielding metric information within near distance”, what (Culham et al., 2008) meant when they suggest that “vergence of the eyes may provide a key signal for encoding near space”, or what Bradshaw et al. (2004) meant when they suggest that “vergence information dominates the control of the transport [reaching] component with minimal contribution from pictorial cues” in reaching and grasping tasks. These are all claims about vergence being a highly effective absolute distance cue in everyday conditions. If vergence is only effective when subjects are consciously attending to their own eye movements, then this literature must be wrong.

Second, conscious awareness of eye movements might contribute to performance in controlled experimental conditions, which is why we controlled for them. But there is no suggestion in the literature that, even in experimental conditions, conscious awareness of eye movements could provide us with the kind of (close to) veridical estimates of absolute distance found in the experimental literature. Could subjects really achieve a relationship of *y* = 0.86*x* + 6.5 (Mon-Williams & Tresilian, 1999) from conscious awareness of eye movements alone?

Third, we know that the visual pathway has access to the vergence signal in LGN, the visual cortex, and the parietal cortex. There is no suggestion that what is being observed in these brain imaging studies is our conscious awareness of our own eye movements. Instead, the suggestion is that the visual system is unconsciously processing the vergence signal, and the question is whether this is actually used to provide distance information. Quinlan & Culham (2007) are not talking about conscious awareness of eye movements when they conclude: “Eye position signals related to the current degree of vergence in dPOS likely supply the dorsal stream with critically important information about object distance with respect to current gaze.”

Fourth, no explanation has been given as to how conscious awareness of eye movements could explain how vergence is supposed to change our visual experience of size (‘size constancy’) or 3D shape (‘depth constancy’). In particular, it becomes very difficult to understand how conscious awareness of eye movements could be involved in disparity scaling. Furthermore, as Regan, Erkelens, & Collewijn (1986) document, changes in size and 3D shape from vergence occur even when there is no appreciable motion-in-depth from vergence (by using large-field stimuli to veto vergence as a motion-in-depth cue).

Fifthly, if subjects are merely responding to a conscious awareness of eye movements, then it is important to recognise this is not visual processing. With the potential exception of blindsight, visual processing affects our visual experience. By contrast, this account of vergence as an absolute distance cue is a purely somatosensory account that has no effect on our visual experience. To give a crude analogy, if I poke you in the eye with a pencil, you may now have a veridical sense of the absolute distance of the pencil, but it would be an aberration of language to call this visual processing. Our point is that under this alternative account, vergence as an absolute distance cue is analogous to the poking in the eye case, rather than the visual processing of absolute distance it was supposed to represent. Elsewhere we have developed this somatosensory account of vergence to encompass not just vergence as an absolute distance cue, but also vergence as a cue to motion-in-depth (Linton, 2018). The key point being that under this account, vergence does not change what we see. We look forward to developing this account to explore, and potentially encompass, directional (version) eye movements as well as depth (vergence) eye movements.

### 4. Change-Blindness

Another suggestion is that the changes in vergence in our experiments were too gradual for the visual system to detect. Similarly, in a series of ‘expanding room’ experiments by Glennerster and collagues, subjects failed to notice gradual changes in vergence and motion parallax: “Subjects seem to ignore information both about vergence angle (to overrule stereopsis) and about stride length (to overrule depth from motion parallax).” (Glennerster, Tcheang, Gilson, Fitzgibbon, & Parker, 2006; see also Rauschecker, Solomon, & Glennerster, 2006; Svarverud, Gilson, & Glennerster, 2010; Svarverud, Gilson, & Glennerster, 2012). But there are two responses to this concern:

First, the failure to notice changes in vergence in the ‘expanding room’ experiments may have little to do with the gradual nature of the vergence change for four reasons: First, the vergence range in those experiments was limited (75cm to 3m). But we know vergence is supposed to be most effective as a distance cue within arm’s reach, and Glennerster et al. (2006) merely interpret their results as indicating that the “efficacy of motion and disparity cues is greater at near viewing distances.” By contrast our experiments test distances within arm’s reach. Second, the ‘expanding room’ experiments use full-field stimuli which we know vetoes motion-in-depth from vergence, even when the change in vergence is far from gradual (up to 13.5°/s in Erkelens & Collewijn, 1985a; 1985b). Third, the pictorial cues in the ‘expanding room’ experiments provide the illusion of a stable scene. So all this demonstrates (as the title of Glennerster et al., 2006 illustrates) is that “humans ignore motion and stereo cues in favor of a fictional stable world”. Finally, Rogers (2011) is highly critical of the ‘expanding room’ experiments being used as evidence of subjects failing to notice gradual vergence changes, and found conflicting results when he tested gradual vergence changes: “the gradualness of the change in interocular separation (and hence vergence demand) did not preclude the appropriate scaling of the disparity-specified ridge surfaces.”

Second, even if the gradual nature of the change is responsible, subjects in the ‘expanding room’ experiments actually notice the change once they have been alerted to its possibility. In this regard the ‘expanding room’ experiments are no different from gradual colour change-blindness experiments where a region of a painting gradually changes in colour without subjects noticing (Simons, Franconeri, & Reimer, 2000; Auvray & O’Regan, 2003). As I’ve already explained in Linton (2017), p.102, this change-blindness is better thought of as cognitive rather than perceptual (can it really be maintained that as a region of the painting changes from red to blue over 30 seconds, the observer’s visual experience remains red over the course of the 30 seconds?). But the important point is this. If you ask subjects in the gradual colour case to discriminate the colour at t_30_ (e.g. by asking them ‘what colour is this region of the picture?’), they can do so accurately even though they don’t detect the change during the experiment. Interestingly, what Glennerster et al. (2006)’s experiment shows is that subjects are actually very good at detecting gradual changes in vergence and motion parallax once they have been alerted to their possibility; i.e. when they re-evaluate the distances in the scene rather than simply assuming the previous depth relations in the scene apply. So in both the gradual colour case, and the gradual vergence and motion parallax case, there is an absolute signal at t_30_ that subjects have access to even if they miss the gradual change from t_1_, t_2_, … t_30_. In conclusion, it would be no criticism of a colour discrimination task that the colours were gradually varied between trials. If the colour was blue at t_1_, and red at t_30_, subjects would still be able to recognise the colour at t_30_ when asked ‘what colour is this region of the picture?’, even though they failed to detect the colour change. So why think the gradual variation of vergence in our distance discrimination experiment should be any different?

### 5. Delta Theta rather than Theta

One suggestion that has been put to us, is that there is a disanalogy between gradual changes in colour, and gradual changes in vergence. This argument suggests that whilst gradual changes in colour have two components (the incremental change in shade from t_1_, t_2_, … t_30_, and the absolute colour at t_30_), with subjects reporting the absolute colour at t_30_, in the case of distance from vergence there is no absolute value at t_30_, only the incremental changes from t_1_, t_2_, … t_30_. Put another way, our experimental results have been interpreted as supporting an intermediate position. We have proved (a) that the visual system is unable to extract absolute distance from static vergence (vergence angle theta), but (b) the visual system may still be able to extract absolute distance from changes in vergence (delta theta), and the reason we don’t detect this ability to extract absolute distance from delta theta is that our vergence changes are too gradual.

The claim of the delta theta account is that small changes in vergence are unconsciously integrated over time to provide us with a measure of absolute vergence. There are five responses to this suggestion:

First, it is a departure from the orthodox interpretation of vergence as an absolute distance cue. See Howard (2008), citing Swenson (1932), Mon-Williams & Tresilian (1999), and Viguier, Clément, & Trotter (2001), that: “Several studies have revealed that … people can judge the absolute distance of a visual target when the only information is provided by static vergence.” Indeed, traditionally the puzzling facet of the literature was that static vergence was such an effective absolute distance cue, but by comparison motion-in-depth from delta theta was not: “the distance of a stationary object can be judged on the basis of vergence alone. So why was motion-in-depth not produced by changing vergence?” (Howard, 2012; motion-in-depth from vergence discussed in Linton, 2018). So it seems surprising that now delta theta is being proposed as the effective absolute distance cue whilst static vergence is not.

Second, why think that the vergence changes in our experiments were too gradual for the visual system to detect? First, we’d have to maintain that the visual system both does have access to these changes in order to make them, but doesn’t have access to them in order to specify absolute distance. Second, these changes are clearly detectable if subjects close one eye (as monocular version eye movements), demonstrating Tyler (1971)’s observation that “two eyes less are less sensitive than one”. So the visual system has access to these independent monocular signals. And all the vergence signal comprises of are these two monocular signals. Third, in order for these two monocular signals not to be experienced as two independent signals, the visual system has to combine them, and in the process of combination supress them (in order to achieve Tyler, 1971’s suppression). But again, this presupposes access to these signals. Fourth, even as binocular eye movements, these eye movements are clearly detectable if they are in the same direction (as binocular version eye movements), rather than opposing directions (as vergence). But as Erkelens & Collewijn (1985a) note, this suggests that the visual system must have access to these binocular eye movements in order to supress them when they are equal and opposite (vergence eye movements), but not when they are equal and in the same direction (version eye movements). To summarise these three points, the visual system clearly has access to these eye movements when they are in the same direction, so why claim that the visual system doesn’t have access to them because the sign for one of the eyes is in the opposite direction?

Third, we have real difficulty making sense of the proposal that the visual system is able to extract absolute distance (theta) purely from a change in vergence (delta theta). As Brenner & van Damme (1998) observe, simply knowing how much the vergence angle has changed “can be of little use for judging distances if we do not know the orientation of the eyes before the change (1 deg of ocular convergence could be due to a shift in gaze from 20 to just over 21 cm or from 2 to approx. 4 m).” Clearly advocates of this position mean to suggest something more than the idea that vergence is a relative depth cue that can be scaled by an independent source of absolute distance information, otherwise every relative depth cue becomes an absolute distance cue by definition. But we really struggle to make sense of what the positive alternative is.

One suggestion is that vergence is combined with Gogel (1969)’s ‘specific distance tendency’, the suggestion that subjects default to a prior of 2-3m. As one reviewer notes, this may relate to ‘dark vergence’ and/or ‘dark accommodation’ (the natural resting state of the eyes), although (a) ‘dark vergence’ and ‘dark accommodation’ tend to be closer (around 1m for vergence, and 76cm for accommodation: see Owens & Liebowitz, 1980; Jaschinski, Jainta, Hoormann, & Walper, 2007), and (b) it is unclear why the visual system should have access to static vergence in the one particular context of ‘dark vergence’, but not more generally.

We should note that this suggestion is a distortion of the traditional relationship between vergence and the ‘specific distance tendency’ posited in the literature (Mon-Williams & Tresilian, 1999). There the suggestion is that vergence is an independent source of absolute distance information whose measurement of absolute distance is tempered by the ‘specific distance tendency’. By contrast, here the suggestion is that ‘specific distance tendency’ usurps vergence as the independent source of our absolute distance information.

In any case there is no evidence for a tendency towards 2-3m (or 1m, or 76cm) in the vergence distance literature (see Fig.1). Mon-Williams & Tresilian (1999) find a slight contraction of the results around 40cm, not the far distances this account would predict. Viguier, Clément, & Trotter (2001) find an underestimation of distances beyond 40cm, not the overestimate that this account suggests. And given (according to this account) the apparent absence of the delta theta in our experiments, we should expect our results to be dominated by the ‘specific distance tendency’. But this doesn’t happen. As Fig.5 demonstrates, the key finding of our results is pervasive variance. There is no sense in which our results cluster around any specific distance. Intriguingly, the same pervasive variance also holds true of Gogel (1969)’s own results, where the standard deviation of distance estimates is the same size as the actual distance estimates themselves. In the presence of such pervasive variance in ours and Gogel (1969)’s results, we conclude that there is no sense in which a ‘specific distance tendency’ meaningfully exists.

Finally, even if the specific distance tendency were to provide the absolute distance for the initial vergence eye movement, each subsequent vergence eye movement would then have to trace its absolute distance back to this initial estimate. We’d be stuck in a near infinite regress trying to integrate over successive eye movements. If you believe that vergence is an important absolute distance cue in everyday viewing, as opposed to single-shot distance estimates in controlled experimental conditions (such as Mon-Williams & Tresilian, 1999; Viguier, Clément, & Trotter, 2001), then this is another serious challenge to the account.

Fourth, as one Reviewer noted, another criticism of this account is that vergence would be de-calibrated by any slow changes in everyday viewing. One response is that vergence could be recalibrated by other absolute distance cues. But it’s important to recognise what this recalibration would entail. Under the delta theta account, the visual system doesn’t know that vergence has gradually changed, so if vergence changes gradually from 20cm to the horizon, the recalibration would have to involve equating the old vergence angle (20cm) with the new viewing distance (the horizon). Presupposing such gross ignorance of the actual vergence state, whilst maintaining an acute awareness in changes in vergence, seems hard to sustain.

Fifth, the argument that the visual system can extract absolute distance (theta) from changes in vergence (delta theta) has been repeatedly put to us as a way of preserving vergence as an important absolute distance cue in spite of our experimental results. But the suggestion that vergence is blind to gradual changes may be just as damaging. If we maintain that vergence is blind to gradual changes, but it later turns out (as one would expect) that subjects are no less accurate in judging near distances in full-cue conditions when the distance of the target is gradually manipulated (e.g. by pointing to objects in full-cue conditions), then advocates of this account would have to concede that there is no benefit from having vergence as an absolute distance cue when reaching for objects; the very scenario where vergence is thought to be at its most important (Loftus, Servos, Goodale, Mendarozqueta, & Mon-Williams, 2004; Melmoth & Grant, 2006; Melmoth, Storoni, Todd, Finlay, & Grant, 2007). So this account risks replacing the ineffectiveness of vergence under my account with the redundancy of vergence under their account.

## Conclusion

Vergence is considered to be one of our most important absolute distance cues. But vergence has never been tested as an absolute distance cue divorced from obvious confounding cues such as binocular disparity. In this article we control for these confounding cues for the first time by gradually manipulating vergence, and find that observers fail to accurately judge distance from vergence. We consider a number of different interpretations of these results. Whilst ad-hoc reinterpretations of vergence as blind to gradual changes, or reliant on delta theta rather than theta, cannot be definitively ruled out, we argue that the most principled response to these results is to question the effectiveness of vergence as an absolute distance cue. Given other absolute distance cues (such as motion parallax and vertical disparities) are limited in application, this poses a real challenge to our contemporary understanding of visual scale (Linton, 2017; Linton, 2018).

## Acknowledgements

This research was conducted under the supervision of Christopher Tyler and Simon Grant. We would also like to thank Matteo Lisi, Joshua Solomon, Michael Morgan, and Chris Hull for their valuable advice, as well as John Barbur for testing the red filters for Experiment 2 in an autorefractor, and Ron Douglas for in testing them in a spectrophotometer. We would also like to thank audiences at the European Conference in Visual Perception (2018), the Center for Visual Science Symposium at the University of Rochester (2018), the Scottish Vision Group (2018), and the Association for the Scientific Study of Consciousness (2019), where this work was presented.

## Open Practice Statement

All data and analysis scripts are accessible in an open access repository: https://osf.io/2xuwn/. Neither of the experiments were formally pre-registered, but the hypothesis that inspired these studies was stated and explored in Linton (2017).

